# Dietary fibers benefits on glucose homeostasis require type 2 conventional dendritic cells in mice fed a high-fat diet

**DOI:** 10.1101/2023.04.19.537402

**Authors:** Adélaïde Gélineau, Geneviève Marcelin, Melissa Ouhachi, Sébastien Dussaud, Lise Voland, Ines Baba, Christine Rouault, Laurent Yvan-Charvet, Karine Clément, Roxane Tussiwand, Thierry Huby, Emmanuel L. Gautier

**Author notes:** Corresponding author **Address correspondence to:** Dr. Emmanuel L. Gautier, INSERM UMR_S 1166, Sorbonne Université, Faculté de Médecine, Hôpital de la Pitié-Salpêtrière, Paris, France.

## Abstract

Diet composition impacts metabolic health and is now recognized to shape the immune system, especially in the intestinal tract. Nutritional imbalance and increased caloric intake are induced by high-fat diet (HFD) in which lipids are enriched at the expense of dietary fibers. Such nutritional challenge alters glucose homeostasis as well as intestinal immunity. Here, we observed that short-term HFD induced dysbiosis, glucose intolerance and decreased intestinal RORγt^+^ CD4 T cells, including peripherally-induced Tregs and IL17-producing (Th17) T cells. However, dietary fiber supplementation of HFD-fed animals was sufficient to maintain RORγt^+^ CD4 T cell subsets and microbial species known to induce them, alongside having a beneficial impact on glucose tolerance. Dietary fiber-mediated normalization of Th17 cells and amelioration of glucose handling required the cDC2 dendritic cell subset in HFD-fed animals, while IL-17 neutralization limited fibers impact on glucose tolerance. Overall, we uncover a novel and pivotal role of cDC2 in the control of the immune and metabolic effects of dietary fibers in the context of HFD feeding.

## INTRODUCTION

Westernized dietary patterns have been associated to the outbreak and sharp increase in noncommunicable diseases (NCDs) over the past decades ^1,2^. Among NCDs, chronic metabolic disorders are naturally influenced by feeding habits. Indeed, while a healthy diet positively impacts host metabolism, changes in the quantity and quality of the dietary intake can favor the onset and persistence of metabolic dysfunctions ^3^. Besides metabolism, the diet is now recognized to impact other organismal functions such as immunity, and this especially holds in the intestinal tract ^4^.

Earlier studies revealed that diets low in vitamins A or aryl hydrocarbon receptor (AhR) ligands alter intestinal lymphocytes homeostasis ^5^. Lately, dietary fibers have received considerable attention with regards to their widespread impact on immune cells and intestinal homeostasis ^6^, adding to their well-known ability to prevent metabolic disorders ^7^. Importantly, fibers are a major source of energy for the intestinal flora, and thus largely contribute to gut microbiota ecology ^8^. Thus, the tightly regulated cross-talk between the gut microbiota and the host immune system is influenced by the dietary intake, which in turn shapes the metabolites, the microbiome community, and the intestinal tract immune subset composition ^9^.

Unbalanced diets such as the high-fat diet (HFD) are usually low in fibers, which promotes the development of metabolic disorders ^10,11^. HFD intake induces alteration across several organs, including the intestine that contributes to systemic metabolic homeostasis by controlling glucose and lipid metabolism ^12–14^. Further, HFD-induced alteration of intestinal immune cells homeostasis concurs to precipitate metabolic dysfunctions ^15,16^. Notably, HFD feeding was shown to alter intestinal lymphocytes by hampering the maintenance of RORγt-expressing CD4 T cells ^17^. Nonetheless, the mechanisms and immune cell subsets that translate dietary cues into intestinal CD4^+^ T cells homeostasis under HFD feeding remain unexplained.

In the intestine, RORγt^+^ CD4 T cells comprise two main populations. It includes regulatory T cells (Tregs) known to be induced peripherally by the microbiota (pTregs) ^18^. These cells are involved in tolerance to the microbiota to avoid type 2 immunity ^19^. The second population consists of IL-17-expressing T helper (Th17) cells known to be induced in response to commensals, especially segmented filamentous bacteria (SFB) ^20^. At the steady state, Th17 cells display a non-inflammatory phenotype and are involved in the regulation of the intestinal barrier ^21–23^.

Here, we report that short-term HFD is sufficient to deregulate the homeostasis of Th17 cells and RORγt^+^ regulatory T cells in the intestinal tract as both subsets decreased. HFD feeding was accompanied by dysbiosis, including a reduction in microbial species known to support RORγt^+^ pTregs and Th17 cells development. Importantly, dietary fiber supplementation prevents the loss of several of these microbial species and is sufficient to preserve both RORγt^+^ pTregs and Th17 cells in HFD-fed animals. We also reveal that dietary fibers require type 2 conventional CD11b^+^ dendritic cells (cDC2) to maintain intestinal Th17 cells, but not RORγt^+^ pTregs, and improve glucose homeostasis in HFD-fed animals. Finally, the impact of dietary fibers on glucose tolerance was limited following IL-17 neutralization. Overall, we were able to functionally link dietary fiber intake to the cDC2 compartment and show that this dendritic cell subset is critical to the control of the immune and metabolic effects of dietary fibers.

## RESULTS

### HFD feeding decreases RORγt^+^ pTregs and Th17 cells in the small intestine and colon

Prolonged high-fat diet (HFD) feeding was previously shown to alter the intestinal immune system ^15,16^. As the gut rapidly adapts to nutritional changes, we asked whether intestinal immune cells are impacted during the first weeks of HFD. Thus, mice were administered either HFD or regular chow diet for 4 weeks. Already at this time point, HFD increased body weight (Fig. 1A), weight gain (Fig. 1B), epidydimal (Fig. 1C), and whole-body fat mass (Fig. 1D). Furthermore, glucose intolerance (Fig. 1E) as well as an elevated HOMA-IR index (Fig. 1F), indicative of impaired glucose control, were already noticed after short-term HFD feeding. Higher HOMA-IR reflected both elevated fasting blood glucose (Fig. 1G) and insulin levels (Fig. 1H). Altogether, short-term HFD feeding altered systemic glucose homeostasis. Morphological changes in the intestinal tract were also evident in HFD-fed animals, including decreased colon weight (Fig. 1I) and length (Fig. 1J), which are commonly associated with altered colonic homeostasis. In addition, and as previously reported ^10^, HFD feeding resulted in decreased caecum weight (Fig. 1K).

**Figure 1.**
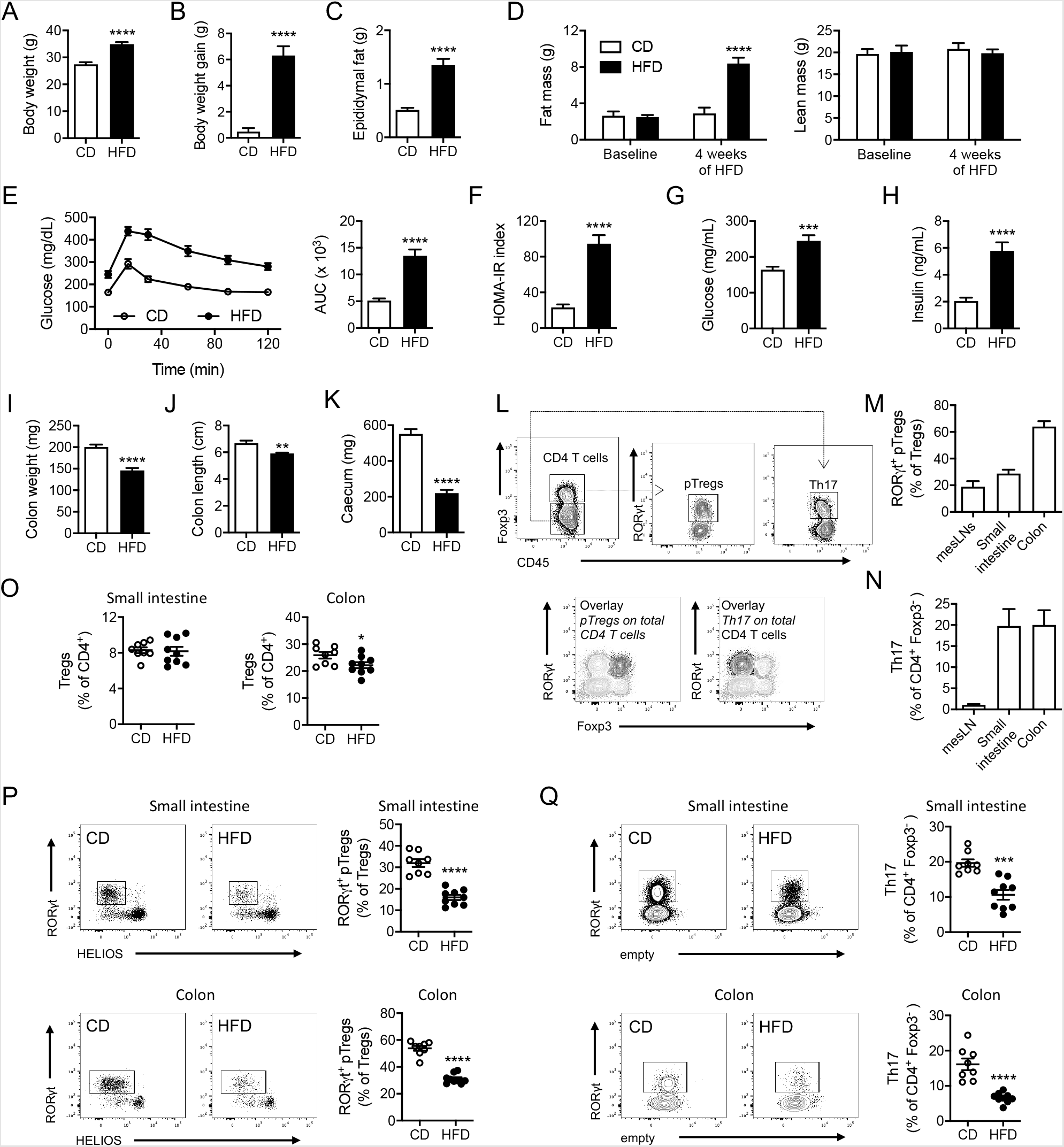
HFD feeding decreases RORγt^+^ pTregs and Th17 cells in the small intestine and colon. **A** to **D**) Body weight (n=13-14 per group) (A), body weight gain (n=13-14 per group) (B), epidydimal fat mass (n=13-14 per group) (C) and body composition (fat and lean mass) (n=5-6 per group) (D) in wild-type mice fed a chow diet (CD) or a high-fat diet (HFD) for 4 weeks. (**E** to **F**) Oral glucose tolerance test and associated area under the curve (AUC) quantification (E), HOMA-IR index measurement (F), plasma glucose (G) and insulin (H) levels in wild-type mice fed a chow diet (CD) or a high-fat diet (HFD) for 4 weeks (n=13-14 per group). (**I** to **K**) Colon weight (n=13-14 per group) (I), colon length (n=9-10 per group) (J) and caecum weight (n=13-14 per group) (K) in wild-type mice fed a chow diet (CD) or a high-fat diet (HFD) for 4 weeks. (**L**) Flow cytometry plots depicting RORγt^+^ CD4 T cells, including RORγt^+^ pTregs and Th17 cells, in the small intestine of chow diet-fed wild-type animals. (**M** and **N**) Flow cytometry analysis of RORγt^+^ pTregs (M) and Th17 cells (N) in the mesenteric lymph nodes (mesLNs) (n=3), the small intestine (n=6) and the colon (n=5) of chow diet-fed wild-type animals. (**O** to **Q**) Flow cytometry analysis of total Tregs (O), RORγt^+^ pTregs (P) and Th17 cells (Q) in the small intestine and colon of wild-type mice fed a chow diet (CD) or a high-fat diet (HFD) for 4 weeks (n=8-9 per group).

We next asked how intestinal lymphocytes were adapting to HFD and focused on RORγt^+^ CD4 T cells, known to be altered under these conditions ^17^. RORγt^+^ CD4 T cells comprise IL-17-producing effector T cells ^24^ and a population of regulatory T cells mostly induced in the periphery (pTregs) ^18,19^ (Fig. 1L), both critical to maintain intestinal homeostasis ^18–20,22,25^. In chow diet-fed animals, the proportion of RORγt^+^ pTregs was similar in the mesenteric lymph nodes (mesLNs) and small intestine, but markedly higher in the colon (Fig. 1M). Th17 effector T cells were more abundant in the small intestine and colonic mucosa as compared to mesLNs, where they are generated (Fig. 1N). In HFD-fed animals, the frequency of total Tregs was unaltered in the small intestine but slightly diminished in the colon (Fig. 1O), while RORγt^+^ pTregs waned at both sites (Fig. 1P). Similarly, RORγt^+^ Th17 cells were also decreased in the small intestinal and colonic lamina propria (Fig. 1Q). Of note, RORγt^+^ CD4 T cell subsets were also reduced when mice were fed the HFD for a longer period of time (14 weeks) (Fig. S1A and S1B). Changes in RORγt^+^ pTregs and Th17 cells could be due to obesity itself, changes in nutrients intake or both. Thus, we studied genetically obese *Ob/Ob* mice maintained on a chow diet or fed with the HFD. Within the small intestine, total Tregs were slightly increased in chow-fed *Ob/Ob* mice, while RORγt^+^ pTregs and Th17 cells were similar to lean, chow-fed controls (Fig. S1C). However, and comparable to WT mice, HFD decreased the frequency of both RORγt^+^ pTregs and Th17 cells in the small intestine (Fig. S1C). In the colon, independently of the genetic background or the diet, total Tregs remained unchanged (Fig. S1D). While colonic RORγt^+^ pTregs and Th17 were significantly reduced in chow-fed obese *Ob/Ob*, HFD further decreased RORγt^+^ pTregs (Fig. S1D). The dysbiosis reported for *Ob/Ob* mice ^26^ likely explains why colonic Th17 cells are already impaired under CD. Overall, these observations suggest that the diet, rather than body weight gain *per se*, has the strongest impact on RORγt^+^ CD4 T cells homeostasis.

Altogether, our observations reveal that short-term HFD negatively impacts systemic glucose homeostasis and profoundly reduces RORγt^+^ pTregs and Th17 cells in the small intestine and colon.

### Dietary fiber supplementation improves glucose tolerance and prevents RORγt^+^ pTregs and Th17 cells loss in HFD-fed animals

Next, we sought to identify the nutritional signals participating in the immune and metabolic alterations induced upon HFD feeding. In the HFD, increased fat content is reached at the expense of cereal starches rich in dietary fibers ^10^. Recent studies pointed out the lack of fermentable dietary fibers as a leading cause of the dysbiosis and metabolic alterations associated with HFD feeding ^10,11,27^. Since dietary fibers were shown to beneficially impact on immune homeostasis and the microbiota ^28^, we wondered whether their administration to HFD-fed animals would be sufficient to preserve RORγt^+^ pTregs and Th17 cells. In order to preserve HFD formulation, we administered fructooligosacharides (FOS) as a fermentable fiber source in the drinking water. Under these conditions, FOS supplementation did not change body weight (Fig. 2A) nor significantly impacted body weight gain (Fig. 2B), epidydimal fat mass (Fig. 2C), body composition (Fig. 2D) and food intake (Fig. 2E). Importantly, glucose tolerance was significantly improved by FOS supplementation (Fig. 2F). This was associated with a lowered HOMA-IR index (Fig. 2G) in FOS-treated animals that reflected decreased circulating insulin levels (Fig. 2H) rather than reduced plasma glucose concentrations (Fig. 2I). In addition, we observed that FOS increased glucose-stimulated insulin secretion (GSIS), suggesting improved pancreatic β-cell function (Fig. 2J). In summary, FOS supplementation improves systemic glucose homeostasis in HFD-fed animals.

**Figure 2.**
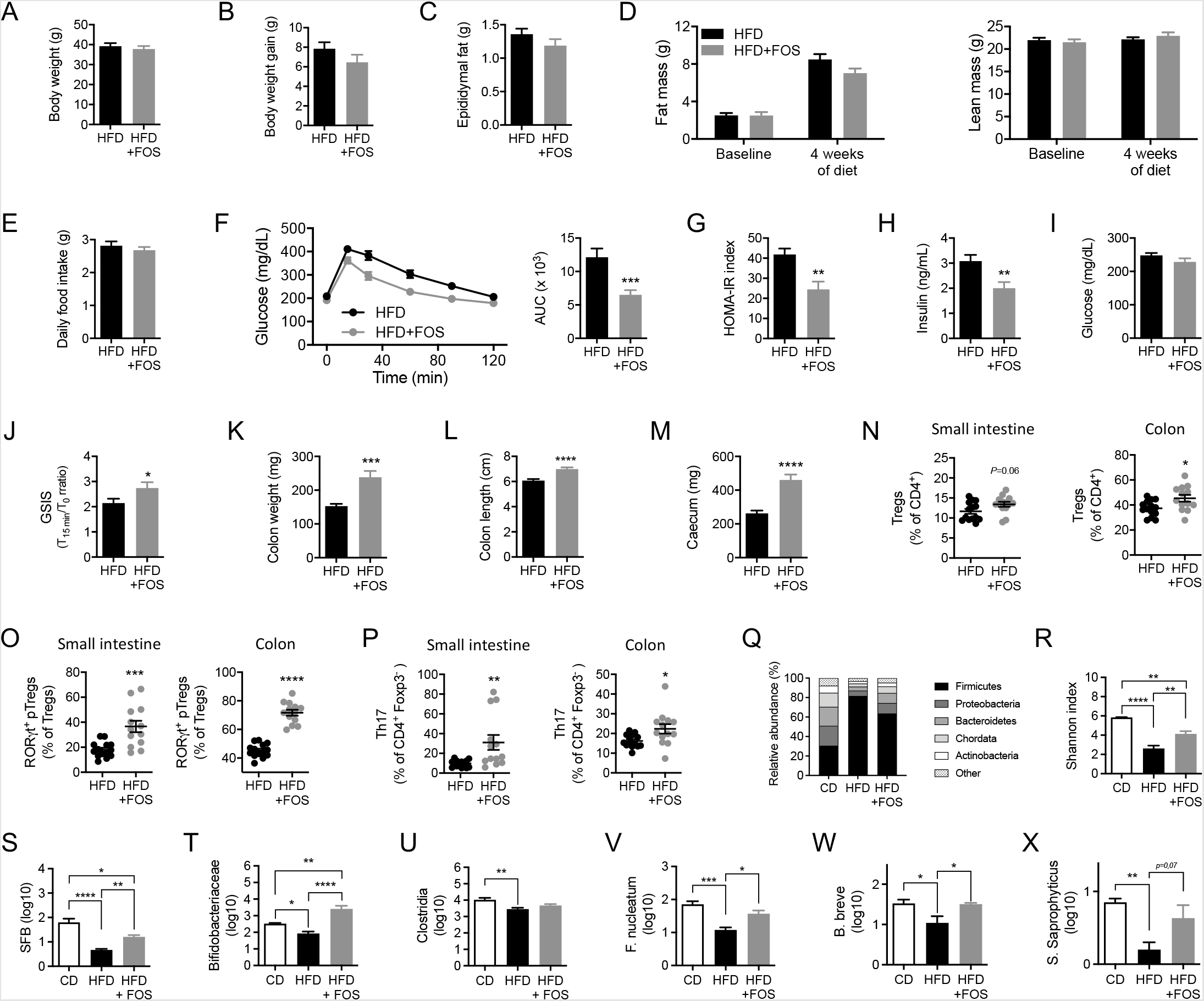
Dietary fiber supplementation improves glucose tolerance and prevents RORγt^+^ pTregs and Th17 cells loss in HFD-fed animals. (**A** to **E**) Body weight (n=15 per group) (A), body weight gain (n=15 per group) (B), epidydimal fat mass (n=14-15 per group) (C), body composition (fat and lean mass) (n=7-10 per group) (D) and daily food intake (n=9 per group) (E) in wild-type mice fed a high-fat diet (HFD) or a high-fat diet supplemented by dietary fibers (fructooligosaccharides, FOS) (HFD+FOS) administered in the drinking water for 4 weeks. (**F** to **J**) Oral glucose tolerance test and associated area under the curve (AUC) quantification (n=15 per group) (F), HOMA-IR index measurement (n=15 per group) (G), plasma insulin levels (n=15 per group) (H), plasma glucose levels (n=15 per group) (I) and glucose-stimulated insulin secretion (GSIS) (n=9-12 per group) (J) in wild-type mice fed a high-fat diet (HFD) or a high-fat diet supplemented with dietary fibers (HFD+FOS) for 4 weeks. (**K** to **M**) Colon weight (n=13-14 per group) (K), colon length (n=9-12 per group) (L) and caecum weight (n=13-14 per group) (M) in wild-type mice fed a high-fat diet (HFD) or a high-fat diet supplemented with dietary fibers (HFD+FOS) for 4 weeks. (**N** to **P**) Flow cytometry analysis of total Tregs (N), RORγt^+^ pTregs (O) and Th17 cells (P) in the small intestine and colon of wild-type mice fed a high-fat diet (HFD) or a high-fat diet supplemented with dietary fibers (HFD+FOS) for 4 weeks (n=13-14 per group). (**Q to X**) Microbiome sequencing and analysis of phylum relative abundance (Q), the Shannon index of microbial diversity (R) and the abundance of specific bacterial class or species including Segmented Filamentous Bacteria (SFB) (S), Bifidobacteria (T), Clostridia (U), F. *nucleatum* (V), B. *breve* (W) and S. *saprophyticus* (X) in wild-type mice fed a chow diet (CD), a high-fat diet (HFD) or a high-fat diet supplemented with dietary fibers (HFD+FOS) for 4 weeks (n=4 per group).

The beneficial effect of FOS supplementation also translated into increased colon weight (Fig. 2K) and length (Fig. 2L) as well as caecum weight (Fig. 2M), reaching values similar to regular chow-fed animals (Fig. 1I-K). Beyond these morphologic parameters, FOS induced a trend towards increased Tregs frequency in the small intestine, which reached statistical significance in the colon (Fig. 2N). More specifically, FOS supplementation prevented the HFD-induced drop in RORγt^+^ pTregs (Fig. 2O) and Th17 cells (Fig. 2P) in the small intestine and colon. Importantly, RORγt^+^ pTregs and Th17 homeostasis were not affected by HFD feeding nor fiber supplementation in other metabolic organs such as the liver (Fig. S2A) and the visceral adipose tissue (Fig. S2B).

Dietary fibers and their deprivation are capital in shaping intestinal bacterial ecology, which intricately dialogs with the immune system ^8^. As RORγt^+^ pTregs and Th17 cells development depends on the microbiota ^18,29^, we asked whether changes in microbial ecology could explain why RORγt^+^ T cell subsets are altered in HFD-fed animals. As expected, the relative abundance of phyla switched towards an increase of the Firmicutes after HFD feeding (Fig. 2Q). As a consequence, microbiota diversity was drastically diminished by the HFD as measured by the Shannon index (Fig. 2T). FOS supplementation was able to partially correct, but not fully restore, Firmicutes over-representation and the loss of microbiota diversity (Fig. 2Q-R). Thus, HFD feeding induced dysbiosis that was partially corrected by fibers supplementation. RORγt^+^ pTregs and Th17 cells are induced by specific genres and taxa of the microbiota ^18,20,30,31^. SFB, the best-known Th17 inducer bacteria^20^, was reduced upon HFD and increased following FOS supplementation (Fig. 2S). At the genre level, we assessed Bifidobacteria prevalence, which is a hallmark of fiber administration ^27,32^ and has been associated with Th17 cells generation ^30^. We found that Bifidobacteria were decreased by the HFD and increased in FOS-supplemented animals (Fig. 2T). On the other hand, the most classically known inducers of RORγt^+^ pTregs are Clostridia strains ^31^. We observed that Clostridia were reduced in HFD-fed mice but fibers did not restore them (Fig. 2U). We also looked at specific strains previously shown to induce the generation of RORγt^+^ pTregs in germ-free mice ^18^. Among them, two strains, F. *nucleatum* and B. *breve*, were reduced by the HFD and significantly increased by fibers (Fig. 2V-W). In addition, S. *Saprophyticus* was also decreased by the HFD and displayed a tendency towards re-induction in FOS-supplemented animals (Fig. 2X). Other RORγt^+^ pTregs inducer strains were not found in our samples (C. *ramosum*, A. *iwoffii*, F. *mortiferum*), unchanged (L. *rhamnosus*) or not relevantly regulated (E. *faecium*, L. *casei*, B. *thetaiotamicron*) (data not shown). Overall, we show that FOS supplementation maintains several bacterial species and genres important for the generation of RORγt^+^ pTregs and/or Th17 cells.

Altogether, we demonstrate that dietary fiber supplementation in HFD-fed animals has a beneficial impact on glucose metabolism, prevents the loss of RORγt^+^ pTregs and Th17 cells and limits the decrease in key microbial species supporting the development of RORγt^+^ CD4 T cell subsets.

### Th17 generation and gut-homing imprinting in the mesenteric lymph nodes are impaired by HFD feeding and corrected upon dietary fiber supplementation

The microbiota data presented above suggests that the decrease in RORγt^+^ CD4 T cells in the intestinal tract of HFD-fed mice could result from their defective priming and development. It could also be potentiated by defective migration to the intestinal tract due to altered gut-homing imprinting. We thus focused on the mesenteric lymph nodes (mesLNs) where RORγt^+^ pTregs and Th17 cells are generated and educated. While total Tregs (Fig. 3A) and RORγt^+^ pTregs (Fig. 3B) were unaltered in the mesLNs of HFD-fed animals, Th17 cells were reduced (Fig. 3C). This revealed that Th17 cells priming was decreased upon HFD. This is consistent with the drop in SFB abundance we observed upon HFD feeding (Fig. 2S). We next evaluated the gut-homing imprinting of the two RORγt^+^ T cell subsets. The frequency of RORγt^+^ pTregs and Th17 cells expressing the gut-homing receptor CCR9 was reduced in HFD-fed mice (Fig. 3D-E). In addition, RORγt^+^ pTregs expressing the gut-homing integrin α4β7 were also decreased in the mesLNs upon HFD (Fig. 3D-E). Together, this indicates that the decline in intestinal Th17 cells observed after HFD feeding is owed to both decreased generation and gut-homing imprinting in the mesLNs, while RORγt^+^ pTregs diminution would solely rely on defective gut-homing imprinting in the mesLN.

**Figure 3.**
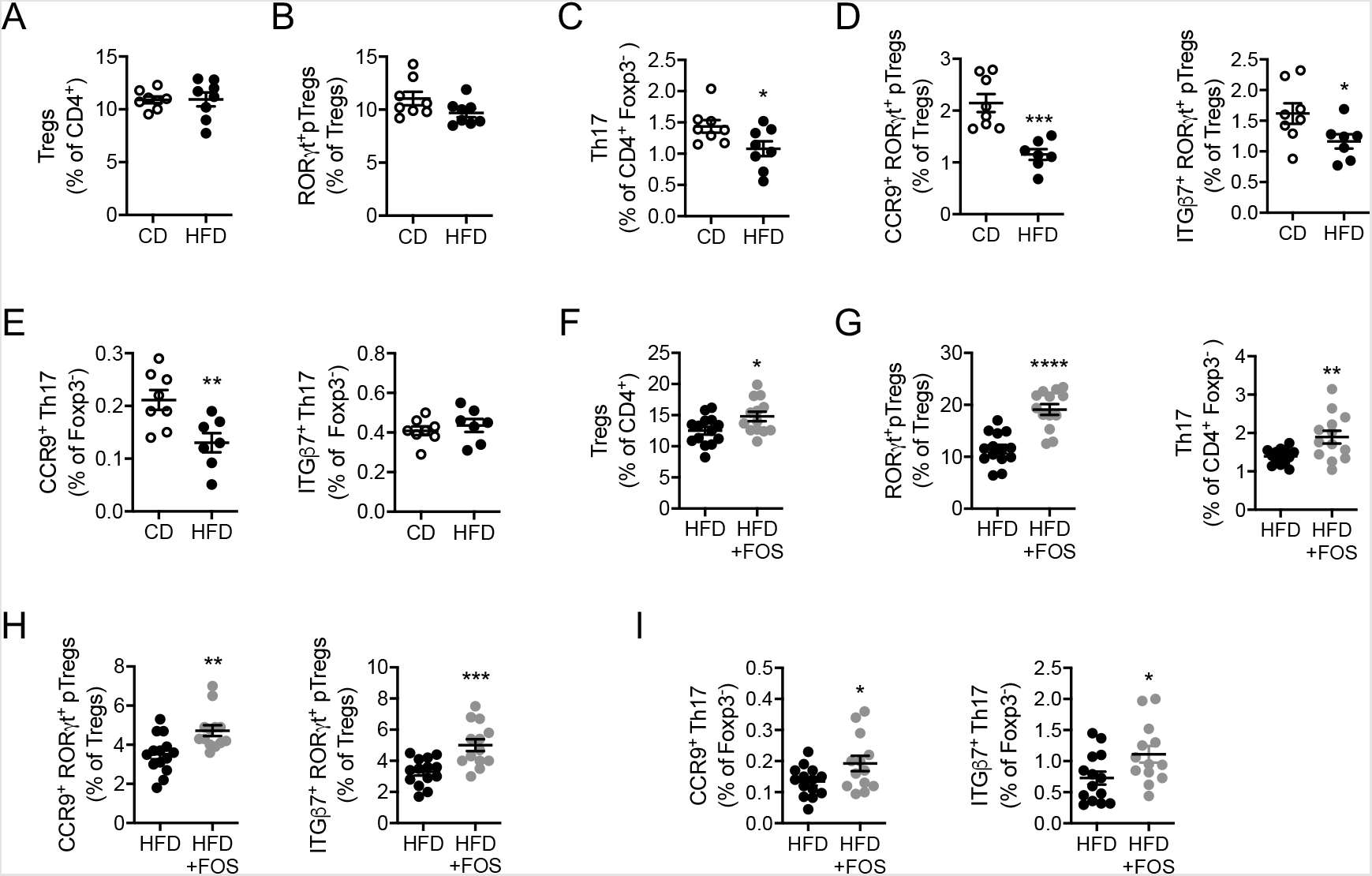
Th17 cells generation and gut-homing imprinting of RORγt^+^ pTregs and Th17 cells are impaired by HFD feeding and corrected upon dietary fiber supplementation. (**A** to **C**) Flow cytometry analysis of total Tregs (A), RORγt^+^ pTregs (B) and Th17 cells (C) in the mesenteric lymph nodes of wild-type mice fed a chow diet (CD) or a high-fat diet (HFD) for 4 weeks (n=8 per group). (**D** and **E**) Flow cytometry analysis of RORγt^+^ pTregs (D) and Th17 cells (E) expressing CCR9 or ITGβ7 in the mesenteric lymph nodes of wild-type mice fed a chow diet (CD) or a high-fat diet (HFD) for 4 weeks (n=7-8 per group). (**F** to **G**) Flow cytometry analysis of total Tregs (F), RORγt^+^ pTregs (G) and Th17 cells (G) in the mesenteric lymph nodes of wild-type mice fed a high-fat diet (HFD) or a high-fat diet supplemented with dietary fibers (HFD+FOS) for 4 weeks (n=13-14 per group). (**H** and **I**) Flow cytometry analysis of CCR9 or ITGβ7-expressing RORγt^+^ pTregs (H) and Th17 cells (I) in the mesenteric lymph nodes of wild-type mice fed a high-fat diet (HFD) or a high-fat diet supplemented with dietary fibers (HFD+FOS) for 4 weeks (n=13-14 per group).

We then sought to decipher whether the fiber supplementation would maintain RORγt^+^ T cells priming and gut-homing imprinting in the mesLNs of HFD-fed animals. FOS supplementation during HFD feeding increased total Tregs (Fig. 3F) as well as RORγt^+^ pTregs and Th17 cells (Fig. 3G) in the mesLNs. In addition, CCR9^+^ and ITGβ7^+^ RORγt^+^ pTregs (Fig. 3H) as well as CCR9^+^ and ITGβ7^+^ Th17 cells (Fig. 3I) were increased after FOS supplementation.

Overall, HFD feeding decreases Th17 cells generation and gut-homing imprinting in the mesLNs, while RORγt^+^ pTregs only show an impairment of their gut-homing imprinting. In this context, FOS supplementation prevents the alterations observed in both RORγt^+^ CD4 T cell subsets.

### IRF4-dependent dendritic cells (cDC2) participate in the homeostasis of RORγt^+^ pTregs and Th17 cells in chow-fed animals

CD103^+^ cDCs control intestinal T cells polarization in the mesLNs ^33^. This DC subset is heterogeneous and can be further subdivided into CD103^+^ CD11b^-^ cDC1 and CD103^+^ CD11b^+^ cDC2 ^34^. CD103^-^ CD11b^+^ cDC2 are also found in the mesLN ^35^ but they remain less well characterized. While cDC1 development depends on the transcription factors BATF3 and IRF8 ^36,37^, cDC2 rely on the transcription factor IRF4 ^38,39^ and were shown to be instrumental in the induction of intestinal Th17 cells ^38,39^. Which DC subset is responsible for the induction of intestinal RORγt^+^ pTregs in adulthood remains, so far, less clear. Given that RORγt^+^ pTregs and Th17 cells share the expression of RORγt and are both altered upon HFD, we wondered if both subsets were primed by cDC2. In chow diet-fed *Itgax*-cre x *Irf4*^flox/flox^ (*Irf4*^ΔDC^) mice, cDC2 were profoundly reduced in the intestinal lamina propria and mesLNs (Fig. S4A-B), as previously reported ^38,39^. CD103^-^ CD11b^+^ cDC2 were slighly reduced in the mesLNs (Fig. S4B) but not the lamina propria (Fig. S4A), while cDC1 were unaltered in the lamina propria (Fig. S4A) and slightly increased in the mesLN (Fig S4B). We found that both Th17 cells (Fig. 4A) and RORγt^+^ pTregs (Fig. 4B) were decreased in the small intestine, colon and mesLNs of *Irf4*^ΔDC^ mice as compared to controls. In addition, the gut-homing imprinting of RORγt^+^ CD4 T cell subsets was decreased in *Irf4*^ΔDC^ (Fig. 4C and 4D). In order to assess whether cDC1 would participate in RORγt^+^ pTregs priming at the steady state, we used chow-fed *Itgax*-cre x *Irf8*^flox/flox^ (*Irf8*^ΔDC^) mice, which lack CD103^+^ CD11b^-^ cDC1 in the intestine (Fig. S5A) and mesLNs (Fig. S5B). In the small intestine, colon and mesLNs, loss of cDC1 had no impact on RORγt^+^ pTregs (Fig. S5C) while Th17 cells were increased (Fig. S5D). Thus, cDC2, but not cDC1, participate to the maintenance of both Th17 cells and RORγt^+^ pTregs.

**Figure 4.**
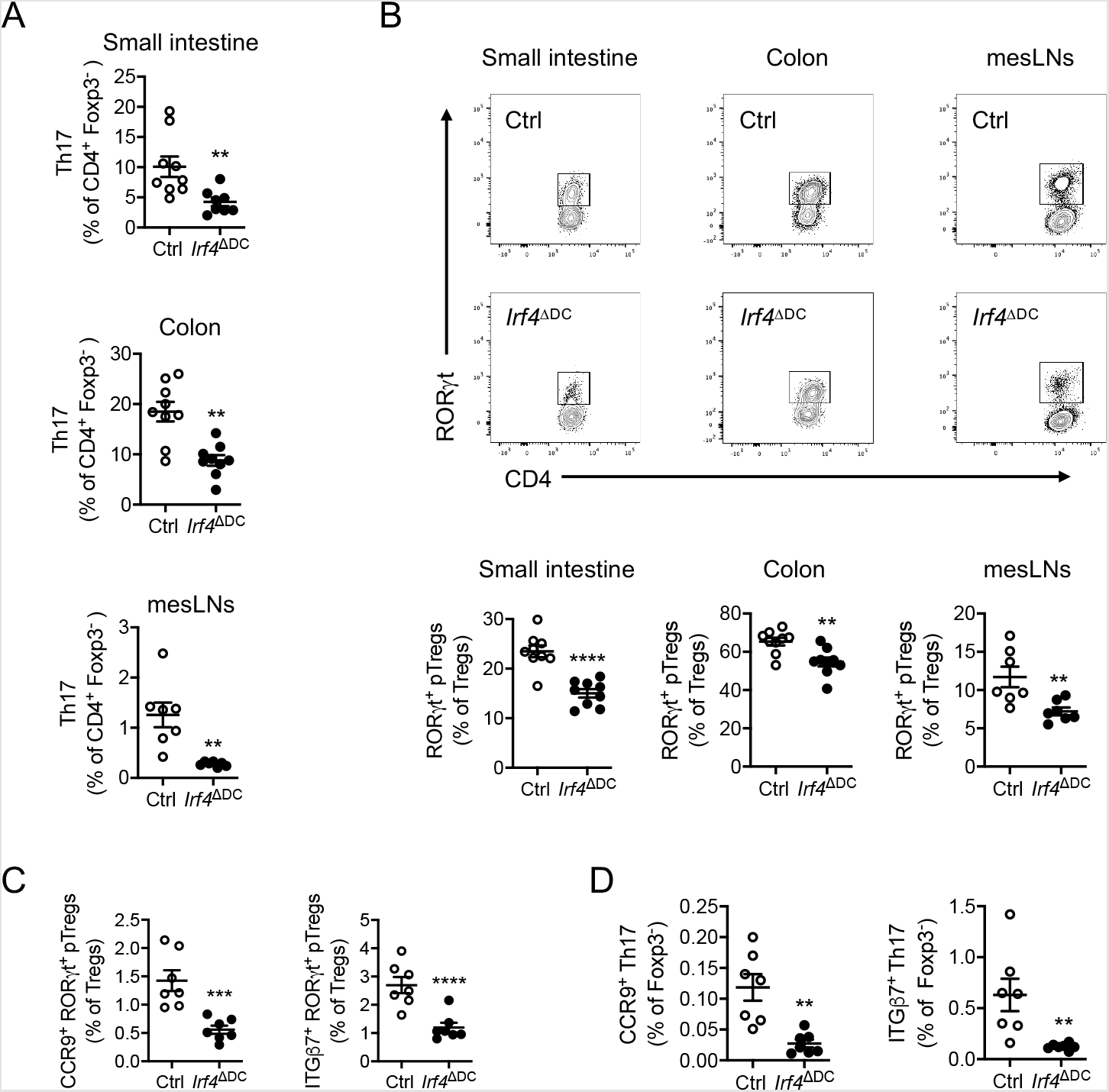
IRF4-dependent dendritic cells (cDC2) participate to the homeostasis of RORγt^+^ pTregs and Th17 cells. (**A**) Flow cytometry analysis of Th17 cells in the small intestine (n=9 per group), colon (n=9 per group) and mesenteric lymph nodes (mesLNs) (n=7 per group) of mice lacking *Irf4* in dendritic cells (*Irf4*^DDC^) and *Irf4*^flox/flox^ controls (ctrl). (**B**) Flow cytometry plots and analysis of RORγt^+^ pTregs in the small intestine (n=9 per group), colon (n=9 per group) and mesenteric lymph nodes (mesLNs) (n=7 per group) of mice lacking *Irf4* in dendritic cells (*Irf4*^DDC^) and *Irf4*^flox/flox^ controls (Ctrl). (**C** and **D**) Flow cytometry analysis of CCR9 or ITGβ7-expressing RORγt^+^ pTregs (C) and Th17 cells (D) in the mesenteric lymph nodes of mice lacking *Irf4* in dendritic cells (*Irf4*^DDC^) and *Irf4*^flox/flox^ controls (Ctrl) (n=7 per group).

We then asked whether cDC subsets were affected upon HFD feeding and dietary fibers supplementation. In HFD-fed animals, we observed a slight decrease in the number of CD103^+^ CD11b^+^ cDC2 in the mesLNs, while CD103^-^ CD11b^+^ cDC2 and CD103^+^ CD11b^-^ cDC1 remained unchanged (Fig. S3B). However, fiber supplementation did not impact on cDC2 numbers in HFD-fed animals (Fig. S3K). This suggests that fiber supplementation prevented HFD-induced loss of RORγt^+^ pTregs and Th17 cells independently from the restoration of cDC2 numbers.Collectively, we show that cDC2 are responsible for the maintenance of both RORγt^+^ CD4 T cell subsets at the steady state, and that cDC2 numbers are not impacted upon fiber supplementation.

### cDC2 control dietary fiber-mediated prevention of Th17 cells loss and glucose tolerance improvement in HFD-fed animals

We showed above that cDC2 numbers are not affected by fibers, but that cDC2 control both Th17 and RORγt^+^ pTregs at the steady state. We also showed that fiber supplementation prevented the HFD-induced decrease in microbial species known to stimulate RORγt^+^ CD4 T cells generation. Thus, we reasoned that cDC2 could control the benefits of dietary fiber by sensing and mediating microbial-derived signals.

To test for the importance of cDC2 in mediating the benefits of FOS supplementation, we fed separated cohorts of *Irf4*^ΔDC^ mice and *Irf4*^flox/flox^ littermate controls with the HFD for 4 weeks and supplemented part of them with FOS. First, we observed that *Irf4*^ΔDC^ and *Irf4*^flox/flox^ control mice responded to FOS supplementation by increasing their colon weight and length as well as caecum weight (Fig. 5A-C) as reported above for wild-type animals (Fig. 2K-M). We next turned our attention to RORγt^+^ CD4 T cell subsets. Unexpectedly, RORγt^+^ pTregs were increased upon FOS supplementation in the small intestine, colon and mesLNs of mice lacking cDC2 to a comparable extent as in *Irf4*^flox/flox^ control animals (Fig. 5D-F). In addition, the lack of cDC2 had no impact on FOS-induced gut-homing imprinting of RORγt^+^ pTregs in the mesLNs (Fig. 5G-H). This suggests that other antigen-presenting cells were capable to compensate for the lack of CD103^+^ CD11b^+^ cDC2 in this context. Nevertheless, while FOS increased Th17 cells in the small intestine, colon and mesLNs of *Irf4*^flox/flox^ control animals, their effect was blunted in *Irf4*^ΔDC^ mice lacking cDC2 (Fig. 5I-K). Moreover, Th17 cells gut-homing imprinting was not improved in FOS-supplemented *Irf4*^ΔDC^ mice as compared to their littermate *Irf4*^flox/flox^ controls (Fig. 5L-M). Thus, while dietary fiber-mediated prevention of RORγt^+^ pTregs loss most likely benefit from cDC2-independent mechanisms, fiber’s impact on Th17 cells was fully dependent on cDC2. We then wondered if the absence of cDC2 had any impact on the beneficial metabolic effects of dietary fiber supplementation in HFD-fed animals. To this aim, separated cohorts of *Irf4*^ΔDC^ mice and *Irf4*^flox/flox^ littermate controls were fed the HFD for 4 weeks and part of the animals were supplemented with FOS. Separated cohorts were studied as we could not perform the metabolic exploration simultaneously with such a number of animals. Body weight (Fig. 6A), weight gain (Fig. 6B), epididymal fat mass (Fig. 6C), body composition (Fig. 6D) and food intake (Fig. 6E) were not significantly modified by FOS supplementation in *Irf4*^ΔDC^ and *Irf4*^flox/flox^ littermate controls. Importantly, while FOS improved glucose tolerance in *Irf4*^flox/flox^ control animals, this effect was abrogated in *Irf4*^ΔDC^ mice lacking cDC2 (Fig. 6F), reflecting the lower HOMA-IR index (Fig. 6G) and better glucose-stimulated insulin secretion (Fig. 6H) of *Irf4*^flox/flox^ animals but not *Irf4*^ΔDC^ mice.

**Figure 5.**
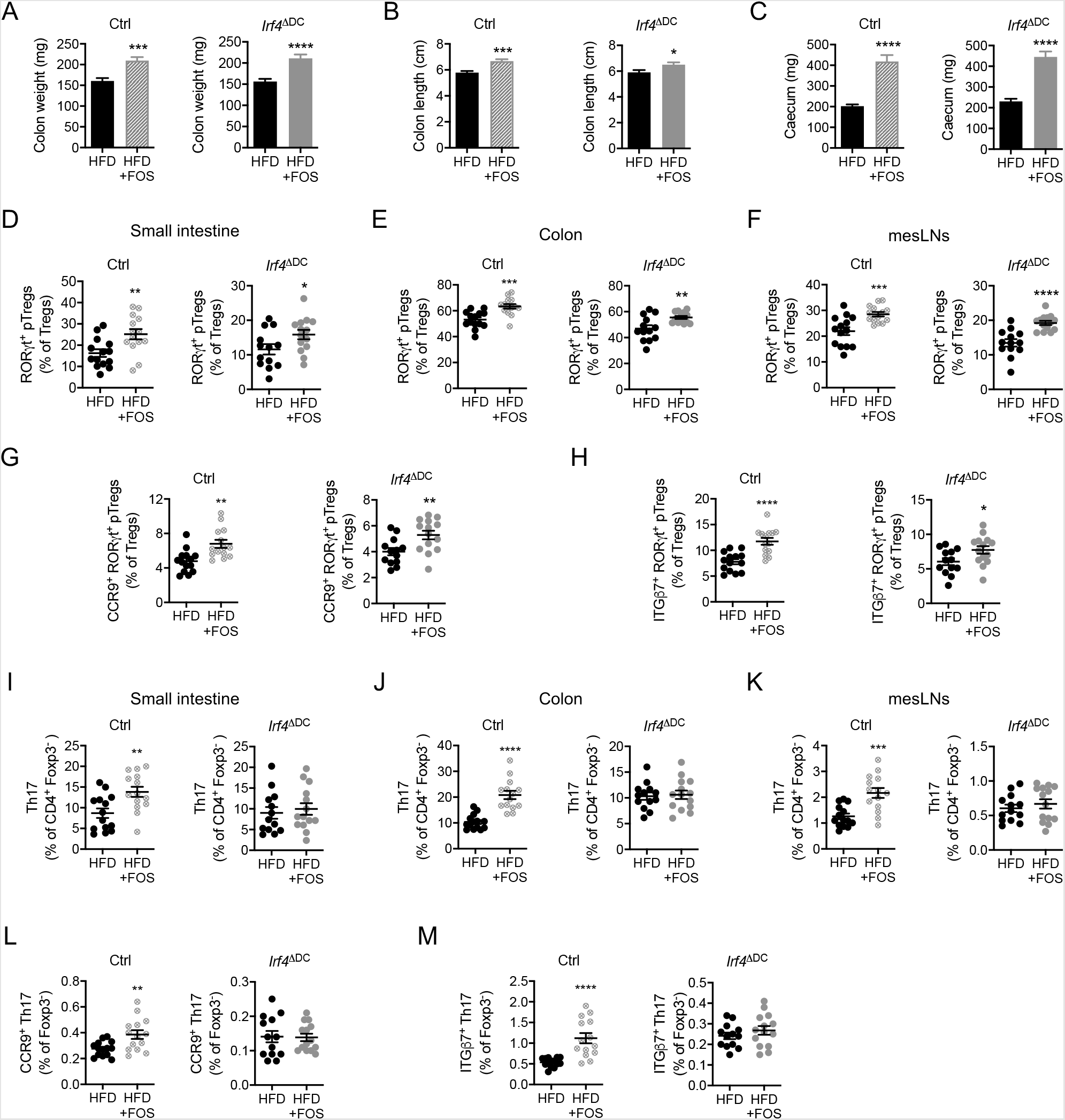
cDC2 control dietary fiber-mediated prevention of Th17 cells loss in HFD-fed animals. (**A** to **C**) Colon weight (A), colon length (B) and caecum weight (C) in mice lacking *Irf4* in dendritic cells (*Irf4*^DDC^) (n=13-14 per group) and *Irf4*^flox/flox^ controls (ctrl) (n=11-14 per group) fed a high-fat diet (HFD) or a high-fat diet supplemented with dietary fibers (HFD+FOS) for 4 weeks. (**D** to **F**) Flow cytometry analysis of RORγt^+^ pTregs in the small intestine (D), colon (E) and mesenteric lymph nodes (mesLNs) (F) of mice lacking *Irf4* in dendritic cells (*Irf4*^DDC^) (n=13-14 per group) and *Irf4*^flox/flox^ controls (ctrl) (n=11-14 per group) fed a high-fat diet (HFD) or a high-fat diet supplemented with dietary fibers (HFD+FOS) for 4 weeks. (**G** and **H**) Flow cytometry analysis of CCR9 (G) or ITGβ7 (H)-expressing RORγt^+^ pTregs in the mesenteric lymph nodes (mesLNs) of mice lacking *Irf4* in dendritic cells (*Irf4*^DDC^) (n=13-14 per group) and *Irf4*^flox/flox^ controls (ctrl) (n=14 per group) fed a high-fat diet (HFD) or a high-fat diet supplemented with dietary fibers (HFD+FOS) for 4 weeks. (**I** to **K**) Flow cytometry analysis of Th17 cells in the small intestine (I), colon (J) and mesenteric lymph nodes (mesLNs) (K) of mice lacking *Irf4* in dendritic cells (*Irf4*^DDC^) (n=13-14 per group) and *Irf4*^flox/flox^ controls (ctrl) (n=14 per group) fed a high-fat diet (HFD) or a high-fat diet supplemented with dietary fibers (HFD+FOS) for 4 weeks. (**L** and **M**) Flow cytometry analysis of CCR9 (L) or ITGβ7 (M)-expressing Th17 cells in the mesenteric lymph nodes (mesLNs) of mice lacking *Irf4* in dendritic cells (*Irf4*^DDC^) (n=13-14 per group) and *Irf4*^flox/flox^ controls (ctrl) (n=14 per group) fed a high-fat diet (HFD) or a high-fat diet supplemented with dietary fibers (HFD+FOS) for 4 weeks.

**Figure 6.**
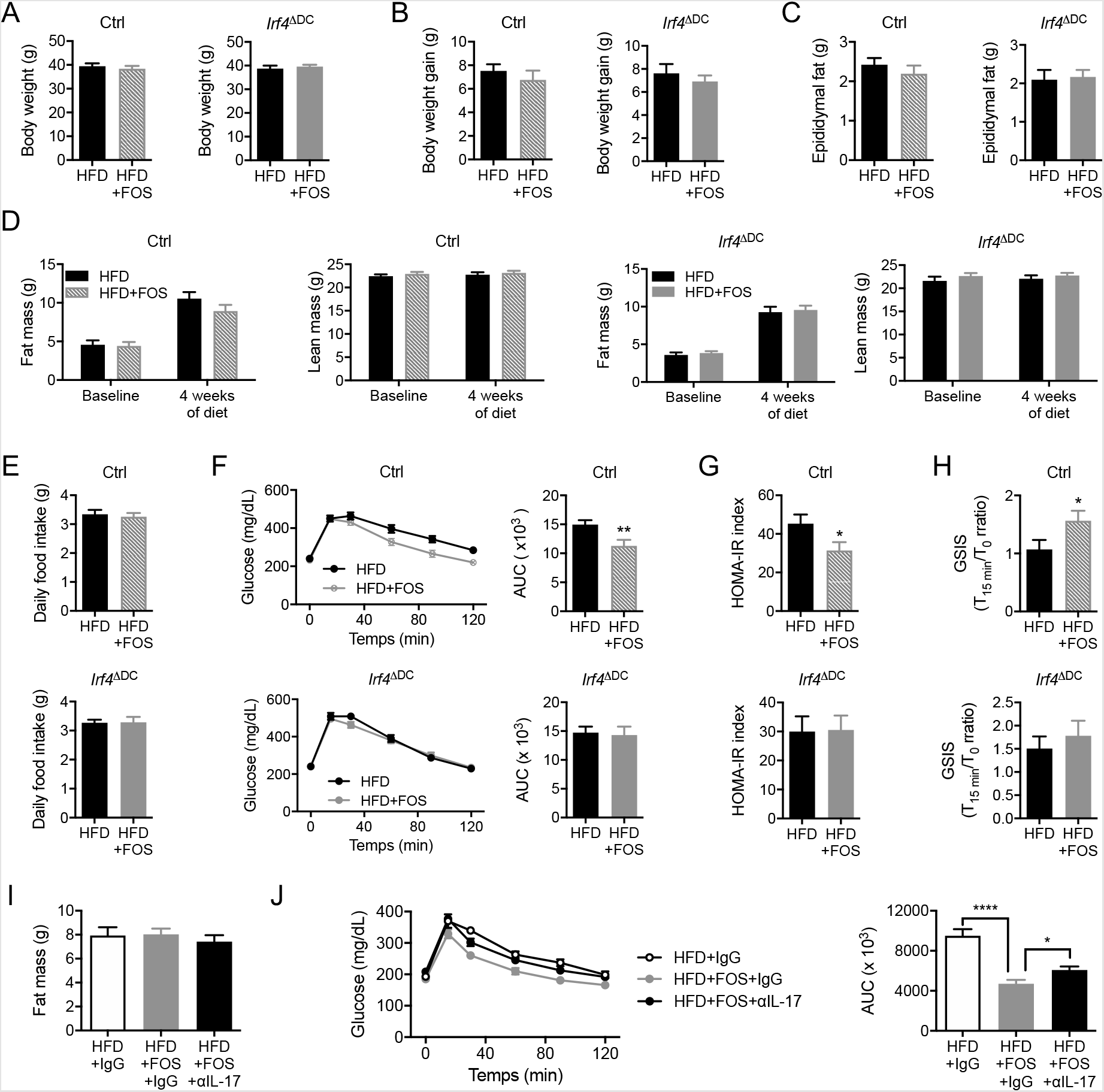
cDC2 control dietary fiber-mediated improvement of glucose tolerance in HFD-fed animals. (**A** to **E**) Body weight (A), body weight gain (B), epidydimal fat mass (C), body composition (fat and lean mass) (D) and daily food intake (E) in *Irf4*^DDC^ mice (*Irf4*^DDC^) (n=6-14 per group) and *Irf4*^flox/flox^ controls (ctrl) (n=11-16 per group) fed a high-fat diet (HFD) or a high-fat diet supplemented with dietary fibers (HFD+FOS) for 4 weeks. (**F** to **H**) Oral glucose tolerance test and associated area under the curve (AUC) quantification (F), HOMA-IR index measurement (G) and glucose-stimulated insulin secretion (GSIS) (H) in *Irf4*^DDC^ mice (*Irf4*^DDC^) (n=8-14 per group) and *Irf4*^flox/flox^ controls (ctrl) (n=12-16 per group) fed a high-fat diet (HFD) or a high-fat diet supplemented with dietary fiber (HFD+FOS) for 4 weeks. (**I** and **J**) Body fat mass (I) and oral glucose tolerance test with the associated area under the curve (AUC) quantification (J) measured after 4 weeks of HFD in mice treated with an isotype control (HFD+IgG, n=6) as well as mice supplemented with dietary fiber and treated with an isotype control (HFD+FOS+IgG, n=9) or antibodies neutralizing IL-17A and IL-17F (HFD+FOS+αIL-17, n=9).

We then asked whether dietary fibers require an intact IL-17 signaling to improve glucose tolerance in HFD-fed mice. To answer that question, we treated HFD-fed animals supplemented with FOS with neutralizing antibodies directed against IL-17A and IL-17F, which are both produced by Th17 cells, or their appropriate isotype control. While IL-17 neutralization did not impact on body fat mass (Fig. 6I), it limited the beneficial impact of FOS supplementation on glucose clearance (Fig. 6J). Thus, the beneficial metabolic impact of dietary fibers partially depends on IL-17 to ameliorate glucose homeostasis, arguing that a cDC2-Th17 axis participates to the dietary fiber-mediated improvement of metabolic fitness in HFD-fed animals.

Together, our work provides new insight regarding the cellular mechanisms by which dietary fibers beneficially impact glucose metabolism. Here, we uncovered a previously unappreciated role for cDC2 in mediating dietary fibers effects on Th17 cells homeostasis and systemic glucose tolerance in the context of HFD feeding.

## DISCUSSION

HFD-induced obesity remains the standard model to study obesity pathogenesis in preclinical models. HFD mimics the human “western” diet, reflecting increased fat content at the expense of dietary fibers. Such nutritional challenge leads to metabolic dysfunctions, including altered glucose homeostasis, and was shown to impact intestinal immune homeostasis. We investigated the effect of a short-term HFD regimen and whether specific changes in intestinal immune subsets participate to the dysregulation of glucose homeostasis. Here, we report that HFD induces dysbiosis, including the reduction in bacterial strains known to favor RORγt^+^ CD4 T cell generation, and decreases intestinal RORγt^+^ pTregs and Th17 cells. Importantly, dietary fiber supplementation was sufficient to prevent the drop in RORγtinducing bacterial species, maintain both RORγt^+^ CD4 T cell subsets and amend glucose homeostasis in HFD-fed animals. The beneficial effect of dietary fibers required cDC2 to preserve Th17 cells and improve glucose tolerance. Overall, our findings unveil a previously unappreciated role of type 2 conventional dendritic cells in mediating the beneficial impact of dietary fiber intake on mucosal immunity and glucose homeostasis in the context of HFD feeding.

RORγt^+^ pTregs and Th17 cells were decreased in the small intestine and colon of HFD-fed animals, as it was previously shown for the whole RORγt^+^ CD4 T cell pool ^17^. This previous observation was attributed to dysbiosis ^17^ as the gut microbiota plays a central role in the induction of RORγt expression in Th17 cells ^29,43^ and peripheral Tregs ^18,19^. Recently, the HFD-mediated changes in gut microbiota ecology were shown to mostly stem from the low dietary fiber content ^11^. We reveal here that dietary fibers limited the HFD-mediated reduction in the Th17-inducing bacteria SFB ^20^ and the RORγt^+^ pTregs-inducing species B. *breve* and F. *nucleatum* ^18^. Dietary fiber supplementation also prevented the decrease of intestinal RORγt^+^ pTregs and Th17 cells in HFD-fed animals, by maintaining their priming and/or gut-homing imprinting in the mesLNs. Overall, we report that nutritional imbalance, *i*.*e*. dietary fiber paucity, triggers dysbiosis and impairs the maintenance of RORγt^+^ pTregs and Th17 cells. A very recent study suggests that high-sugar content can also lead to reduced SFB abundance and intestinal Th17 cells upon HFD feeding ^44^. This sugar-induced effect is dependent upon competition mechanisms with other members of the microbiota. Thus, different nutritional cues, such as the paucity of readily fermentable dietary fibers or an excess of sugar in the diet, favor an imbalance in microbiota species including decreased SFB abundance.

The beneficial effect of dietary fibers in restoring RORγt^+^ CD4 T cell subsets homeostasis likely relied on cDCs as both RORγt^+^ pTregs and Th17 cells need cDCs to develop ^19,38,39^. First, consistent with the previously described role of *Irf4*-dependent cDC2 in Th17 cells generation ^38,39^, we found that fiber-mediated preservation of Th17 cells in HFD-fed animals required cDC2. Then, we observed that cDC2, but not cDC1, participated to RORγt^+^ pTregs priming at the steady state. Yet, dietary fiber supplementation led to the maintenance of RORγt^+^ pTregs in HFD-fed cDC2-deficient animals. Even though cDC2 have a dominant impact on RORγt^+^ pTregs at the steady state, a significant proportion of RORγt^+^ pTregs remained in their absence. This suggests that cDC1 could somewhat compensate the absence of cDC2 at the steady state and explain why RORγt^+^ pTregs were retained upon fiber supplementation in HFD-fed cDC2-deficient animals. Plasmacytoid DCs (pDCs) could also play a role in RORγt^+^ pTregs homeostasis. Indeed, both cDC1, cDC2 and pDCs are lacking in *Cd11c*-cre x LsL-ROSA-DTA mice ^45^ in which RORγt^+^ pTregs do not develop ^19^, and pDCs were previously shown to favor RORγt^+^ pTreg generation in the intestinal tract ^46^. A recent study suggests that different intestinal DC subsets, including CD103^+^ cDC1 and cDC2 as well as CD103^-^ CD11b^+^ cDCs, have the ability to induce RORγt^+^ pTregs generation depending on the context ^47^. Finally, other cell types such as ILC3 ^48^ or RORγt^+^ antigen-presenting cells ^49,50^ may also be involved in this compensation. In summary, while dietary fiber intake maintains RORγt^+^ pTregs homeostasis in the absence of cDC2, Th17 cells observe a strict dependency on cDC2 to benefit from fiber supplementation.

Since dietary fibers did not fully maintain RORγt^+^ CD4 T cells in the absence of cDC2, we tested whether cDC2 deficiency altered fiber’s ability to improve glucose handling. We observed that dietary fibers failed to improve glucose tolerance in HFD-fed mice lacking cDC2. We thus identified a key role for cDC2 in mediating the beneficial metabolic effects of dietary fibers in HFD-fed animals. This effect did not depend on RORγt^+^ pTregs but partially relied on Th17 cells since they were not restored in dietary fiber-treated cDC2-deficient animals and IL-17 neutralization limited the full impact of dietary fiber on glucose tolerance. On their side, RORγt^+^ pTregs might control other aspects of intestinal homeostasis given their ability to prevent Th2-driven intestinal inflammation ^19^. Importantly, while we focused our attention on particular subsets of CD4 T cells, other alterations in immune intestinal cells have been reported upon long-term HFD feeding ^15,16^, including an increase in Th1 IFNγ-producing CD4 T cells ^51^. Since intestinal Th1 responses are controlled by cDC1 ^52^, further work would be needed to assess the role of cDC1 in HFD-induced metabolic alterations. Together, we show cDC2 are critical to mediate the beneficial impact of dietary fibers on Th17 cells homeostasis and glucose homeostasis in HFD-fed animals.

Dietary fibers were previously shown to protect against HFD-induced obesity. In these studies, adding fibers to the diet markedly limited weight gain and fat mass expansion, leading to an improvement in glucose tolerance ^10,27^. Here, we delivered fibers in the drinking water to keep the diet formulation similar between groups. Under our experimental conditions, fibers were capable to improve glucose handling independently from any major effect on adiposity. Improved glucose tolerance appeared independent from glucose absorption since blood glucose levels peaked similarly in fiber-treated animals and controls after oral glucose challenge. However, we observed that fiber supplementation accelerates glucose clearance together with improved glucose-stimulated insulin secretion. In this context, multiple mechanisms may concur to ameliorate glucose tolerance in fiber-treated animals.

In summary, the lack of dietary fiber is the nutritional cue leading to decreased intestinal RORγt^+^ pTregs and Th17 cells in HFD-fed animals while cDC2 link dietary fiber intake to Th17 homeostasis and the improvement of glucose tolerance. Overall, we provide new insight regarding the cellular mechanisms by which dietary fibers beneficially impact glucose metabolism. More specifically, we uncover a novel and pivotal role of cDC2 in the control of the immune and metabolic effects of dietary fibers.

## METHODS

### Mice and housing

Mice were housed in individually ventilated cages at a temperature of 22°C and maintained under specific pathogen-free conditions on a 12-hour light and dark cycle with ad libitum access to water and diet (A04; Safe-Diets). Age-matched male mice were grouped by cages at weaning according to their genotype.

Wild-type C57BL/6J mice were from Charles River and bred in-house. *Ob/+* (B6.Cg-*Lep*^ob^/J) mice were from Charles River and bred in house to generate obese *Ob/Ob* mice and lean littermate controls (including *Ob/+* and *+/+* animals). *Itgax*-cre (B6.Cg-Tg(Itgax-cre)1-1Reiz/J), *Irf4*^flox/flox^ (B6.129S1-*Irf4*^*tm1Rdf*^/J) and *Irf8*^flox/flox^ (B6(Cg)-*Irf8*^*tm1*.*1Hm*^/J) were all obtained from the Jackson Laboratory. *Itgax*-cre mice were crossed to *Irf4*^flox/flox^ animals in our facility, while *Itgax*-cre x *Irf8*^flox/flox^ were directly imported from the Tussiwand lab (Basel Institute, Switzerland). Littermate cre-negative mice were used as controls and germline deletion events were screened as previously described ^38,52^.

All animal procedures were in accordance with the Guide for the Care and Use of Laboratory Animals published by the European Commission Directive 86/609/EEC and given authorization from the French Ministry of Research.

### Diet and treatment

For high-fat diet (HFD)-induced metabolic dysfunctions studies, male mice were fed a HFD in which 60% of kilocalories come from fat (D12492, Research Diets) and were compared to chow diet-fed animals (A04; Safe-Lab). The duration of feeding was indicated in the text and figure legends. For dietary fiber supplementation studies, mice were given fructooligosaccharides (obtained from Sigma-Aldrich or Beneo) diluted at 7.5% in filtered drinking water. The solution was renewed every 2 to 3 days for the entire diet duration. IL-17A (clone 17F3) and IL17F (clone MM17F8F5.1A9) neutralizing antibodies, as well as their appropriate isotype control (MOPC-21), were obtained from BioXcell. Antibodies (200 μg per injection) were injected intraperitoneally 3 times a week over the 4 weeks period of HFD feeding.

### Glucose metabolism assessment

For assessment of oral glucose tolerance, mice were fasted for 5 hours prior to glucose intragastric gavage at a dose of 1.5 grams per kg of body weight. Glycaemia was measured with a glucometer (Accu-Check, Roche) at baseline and 15, 30, 60, 90 and 120 minutes after gavage. Blood was also collected at baseline and 15 min after gavage for insulin dosage. Insulin dosage was performed with the mouse ultrasensitive insulin ELISA kit from Alpco. The glucose-stimulation insulin secretion index was calculated as the ratio of blood insulin levels measured 15 min after glucose gavage to blood insulin levels at baseline. Finally, the HOMA-IR index was calculated with the following formula: fasting plasma insulin (mU/mL) × fasting plasma glucose (mm/L)/22.5.

### Fat and lean mass measurement

Fat and lean mass were measured by TD-NMR using a MinispecPlus LFII90 body composition analyzer (Bruker; PreclinICAN Plateform, Paris).

### Tissue processing and cell suspension preparation

For isolation of lamina propria leucocytes, freshly harvested intestines and colons were quickly washed in PBS, opened and cut into smaller pieces. To remove epithelial cells, samples were placed into 40 mL of PBS (no calcium and magnesium) containing glucose (1g/L), HEPES (10mM), EDTA (5mM), fetal bovine serum (5%) and dithiothreitol (0.5%), and incubated for 30 min at 37°C under vigorous agitation. After cells were washed 5 times in 40 mL PBS, samples were chopped with scissors and placed in the digestion solution. The digestion solution consisted of HBSS (with calcium and magnesium) containing fetal bovine serum (3%), collagenase D (1.25 mg/mL, Sigma-Aldrich), DNase (10 U/mL, Sigma-Aldrich). Digestion was performed at 37°C under agitation for 30 min. After completion, cells suspensions were passed through a 18G needle before filtration on a 70μm filter, washed and finally resuspended in PBS containing BSA (1%).

Pooled mesenteric lymph nodes were cut open with a needle and digested as described above. Cell suspensions were eventually resuspended in PBS containing BSA (1%) before staining.

### Flow cytometry

Antibodies were purchased from BioLegend, ThermoFisher Scientific and BD Biosciences. The following markers and clones were used: CD11c (N418), MHC-II (I-A/I-E, M5/114.15.2), CD103 (2E7), CD11b (M1/70), RORγt (Q31-378), Foxp3 (FJK-16s), CD4 (GK1.5), CCR9 (CW-1.2), ITGβ7 (DATK32), CD64 (X54-5/7.1) and CD45 (30-F11). Cell suspensions were stained with appropriate antibodies for 30 min on ice. Intracellular staining was performed using the Foxp3 staining kit from ThermoFisher Scientific.

Data were acquired on a BD LSRFortessa™ flow cytometer (BD Biosciences) and analyzed with FlowJo software (Tree Star).

### Microbiome sequencing and analysis

Fecal DNA was extracted using the NucleoMag DNA Microbiome kit (Macherey-Nagel) and sequenced using the MinION from Oxford Nanopore Technologies (ONT). The DNA library was prepared with the Ligation Sequencing Kit with multiplexing (ONT). Sequencing analysis was performed as previously described ^53^. Sequencing data are available upon request.

### Quantification and statistical analysis

Statistical significance of differences was performed using GraphPad Prism (GraphPad Software). Two-tailed Student’s t-test was used to assess the statistical significance of the difference between means of two groups. Graphs depicted the mean ± SEM. Statistical significance is represented as follows: *P<0.05, **P<0.01, ***P<0.001 and ****P<0.0001.

## ACKNOWLEDGMENTS

This work was supported by grants to ELG from the Fondation de France (project number 00056835), the Agence Nationale pour la Recherche (ANR-17-CE14-0009, ANR-17-CE14-0023 and ANR-21-CE14-0023) and from the city of Paris (Emergence-s-program). Melissa Ouhachi received a one-year doctoral fellowship from the Nouvelle Société Française d’Athérosclérose (NSFA).

## AUTHOR CONTRIBUTIONS

A.G. provided intellectual input, designed and performed experiments, analyzed and interpreted data, and wrote the manuscript. G.M., provided intellectual input, designed and performed experiments, analyzed and interpreted data, and edited the manuscript. M.O., I.B., C.R., S.D. and L.V., performed experiments and analyzed data. L.Y.C, R.T and K.C., provided intellectual input and edited the manuscript. T.H., provided intellectual input, designed experiments and edited the manuscript. E.L.G., conceptualized and supervised the study, designed experiments, analyzed and interpreted data, and wrote the manuscript.

## REFERENCES

1. Ezzati, M. & Riboli, E. Behavioral and dietary risk factors for noncommunicable diseases. N. Engl. J. Med. 369, 954–964 (2013).

2. GBD 2013 Risk Factors Collaborators et al. Global, regional, and national comparative risk assessment of 79 behavioural, environmental and occupational, and metabolic risks or clusters of risks in 188 countries, 1990-2013: a systematic analysis for the Global Burden of Disease Study 2013. Lancet Lond. Engl. 386, 2287–2323 (2015).

3. Christ, A., Lauterbach, M. & Latz, E. Western Diet and the Immune System: An Inflammatory Connection. Immunity 51, 794–811 (2019).

4. Thorburn, A. N., Macia, L. & Mackay, C. R. Diet, metabolites, and ‘western-lifestyle’ inflammatory diseases. Immunity 40, 833–842 (2014).

5. Veldhoen, M. & Brucklacher-Waldert, V. Dietary influences on intestinal immunity. Nat. Rev. Immunol. 12, 696–708 (2012).

6. Daïen, C. I., Pinget, G. V., Tan, J. K. & Macia, L. Detrimental Impact of Microbiota-Accessible Carbohydrate-Deprived Diet on Gut and Immune Homeostasis: An Overview. Front. Immunol. 8, 548 (2017).

7. Reynolds, A. et al. Carbohydrate quality and human health: a series of systematic reviews and meta-analyses. Lancet Lond. Engl. 393, 434–445 (2019).

8. Makki, K., Deehan, E. C., Walter, J. & Bäckhed, F. The Impact of Dietary Fiber on Gut Microbiota in Host Health and Disease. Cell Host Microbe 23, 705–715 (2018).

9. Levy, M., Thaiss, C. A. & Elinav, E. Metabolites: messengers between the microbiota and the immune system. Genes Dev. 30, 1589–1597 (2016).

10. Chassaing, B. et al. Lack of soluble fiber drives diet-induced adiposity in mice. Am. J. Physiol. Gastrointest. Liver Physiol. 309, G528–541 (2015).

11. Dalby, M. J., Ross, A. W., Walker, A. W. & Morgan, P. J. Dietary Uncoupling of Gut Microbiota and Energy Harvesting from Obesity and Glucose Tolerance in Mice. Cell Rep. 21, 1521–1533 (2017).

12. Drucker, D. J. The role of gut hormones in glucose homeostasis. J. Clin. Invest. 117, 24–32 (2007).

13. Gribble, F. M. The gut endocrine system as a coordinator of postprandial nutrient homoeostasis. Proc. Nutr. Soc. 71, 456–462 (2012).

14. Ko, C.-W., Qu, J., Black, D. D. & Tso, P. Regulation of intestinal lipid metabolism: current concepts and relevance to disease. Nat. Rev. Gastroenterol. Hepatol. 17, 169–183 (2020).

15. Winer, D. A., Luck, H., Tsai, S. & Winer, S. The Intestinal Immune System in Obesity and Insulin Resistance. Cell Metab. 23, 413–426 (2016).

16. Winer, D. A., Winer, S., Dranse, H. J. & Lam, T. K. T. Immunologic impact of the intestine in metabolic disease. J. Clin. Invest. 127, 33–42 (2017).

17. Garidou, L. et al. The Gut Microbiota Regulates Intestinal CD4 T Cells Expressing RORγt and Controls Metabolic Disease. Cell Metab. 22, 100–112 (2015).

18. Sefik, E. et al. MUCOSAL IMMUNOLOGY. Individual intestinal symbionts induce a distinct population of RORγ+ regulatory T cells. Science 349, 993–997 (2015).

19. Ohnmacht, C. et al. MUCOSAL IMMUNOLOGY. The microbiota regulates type 2 immunity through RORγt+ T cells. Science 349, 989–993 (2015).

20. Ivanov, I. I. et al. Induction of intestinal Th17 cells by segmented filamentous bacteria. Cell 139, 485–498 (2009).

21. Omenetti, S. et al. The Intestine Harbors Functionally Distinct Homeostatic Tissue-Resident and Inflammatory Th17 Cells. Immunity 51, 77–89.e6 (2019).

22. Chung, H. et al. Gut immune maturation depends on colonization with a host-specific microbiota. Cell 149, 1578–1593 (2012).

23. Lécuyer, E. et al. Segmented filamentous bacterium uses secondary and tertiary lymphoid tissues to induce gut IgA and specific T helper 17 cell responses. Immunity 40, 608–620 (2014).

24. Ivanov, I. I. et al. The orphan nuclear receptor RORgammat directs the differentiation program of proinflammatory IL-17+ T helper cells. Cell 126, 1121–1133 (2006).

25. Conti, H. R. et al. Th17 cells and IL-17 receptor signaling are essential for mucosal host defense against oral candidiasis. J. Exp. Med. 206, 299–311 (2009).

26. Ley, R. E. et al. Obesity alters gut microbial ecology. Proc. Natl. Acad. Sci. U. S. A. 102, 11070–11075 (2005).

27. Zou, J. et al. Fiber-Mediated Nourishment of Gut Microbiota Protects against Diet-Induced Obesity by Restoring IL-22-Mediated Colonic Health. Cell Host Microbe 23, 41–53.e4 (2018).

28. Tan, J. K., McKenzie, C., Mariño, E., Macia, L. & Mackay, C. R. Metabolite-Sensing G Protein-Coupled Receptors-Facilitators of Diet-Related Immune Regulation. Annu. Rev. Immunol. 35, 371–402 (2017).

29. Ivanov, I. I. et al. Specific microbiota direct the differentiation of IL-17-producing T-helper cells in the mucosa of the small intestine. Cell Host Microbe 4, 337–349 (2008).

30. Tan, T. G. et al. Identifying species of symbiont bacteria from the human gut that, alone, can induce intestinal Th17 cells in mice. Proc. Natl. Acad. Sci. U. S. A. 113, E8141–E8150 (2016).

31. Atarashi, K. et al. Induction of colonic regulatory T cells by indigenous Clostridium species. Science 331, 337–341 (2011).

32. Gibson, G. R., Beatty, E. R., Wang, X. & Cummings, J. H. Selective stimulation of bifidobacteria in the human colon by oligofructose and inulin. Gastroenterology 108, 975–982 (1995).

33. Coombes, J. L. & Powrie, F. Dendritic cells in intestinal immune regulation. Nat. Rev. Immunol. 8, 435–446 (2008).

34. Merad, M., Sathe, P., Helft, J., Miller, J. & Mortha, A. The dendritic cell lineage: ontogeny and function of dendritic cells and their subsets in the steady state and the inflamed setting. Annu. Rev. Immunol. 31, 563–604 (2013).

35. Cerovic, V. et al. Intestinal CD103(-) dendritic cells migrate in lymph and prime effector T cells. Mucosal Immunol. 6, 104–113 (2013).

36. Edelson, B. T. et al. Peripheral CD103+ dendritic cells form a unified subset developmentally related to CD8alpha+ conventional dendritic cells. J. Exp. Med. 207, 823– 836 (2010).

37. Ginhoux, F. et al. The origin and development of nonlymphoid tissue CD103+ DCs. J. Exp. Med. 206, 3115–3130 (2009).

38. Persson, E. K. et al. IRF4 transcription-factor-dependent CD103(+)CD11b(+) dendritic cells drive mucosal T helper 17 cell differentiation. Immunity 38, 958–969 (2013).

39. Schlitzer, A. et al. IRF4 transcription factor-dependent CD11b+ dendritic cells in human and mouse control mucosal IL-17 cytokine responses. Immunity 38, 970–983 (2013).

40. Chandarana, K. et al. Peripheral activation of the Y2-receptor promotes secretion of GLP-1 and improves glucose tolerance. Mol. Metab. 2, 142–152 (2013).

41. Cani, P. D. et al. Changes in gut microbiota control metabolic endotoxemia-induced inflammation in high-fat diet-induced obesity and diabetes in mice. Diabetes 57, 1470– 1481 (2008).

42. Cani, P. D. et al. Metabolic endotoxemia initiates obesity and insulin resistance. Diabetes 56, 1761–1772 (2007).

43. Atarashi, K. et al. ATP drives lamina propria T(H)17 cell differentiation. Nature 455, 808– 812 (2008).

44. Kawano, Y. et al. Microbiota imbalance induced by dietary sugar disrupts immunemediated protection from metabolic syndrome. Cell 185, 3501–3519.e20 (2022).

45. Ohnmacht, C. et al. Constitutive ablation of dendritic cells breaks self-tolerance of CD4 T cells and results in spontaneous fatal autoimmunity. J. Exp. Med. 206, 549–559 (2009).

46. Uto, T. et al. Critical role of plasmacytoid dendritic cells in induction of oral tolerance. J. Allergy Clin. Immunol. 141, 2156–2167.e9 (2018).

47. Russler-Germain, E. V. et al. Gut Helicobacter presentation by multiple dendritic cell subsets enables context-specific regulatory T cell generation. eLife 10, e54792 (2021).

48. Lyu, M. et al. ILC3s select microbiota-specific regulatory T cells to establish tolerance in the gut. Nature 610, 744–751 (2022).

49. Kedmi, R. et al. A RORγt+ cell instructs gut microbiota-specific Treg cell differentiation. Nature 610, 737–743 (2022).

50. Akagbosu, B. et al. Novel antigen-presenting cell imparts Treg-dependent tolerance to gut microbiota. Nature 610, 752–760 (2022).

51. Luck, H. et al. Regulation of obesity-related insulin resistance with gut anti-inflammatory agents. Cell Metab. 21, 527–542 (2015).

52. Luda, K. M. et al. IRF8 Transcription-Factor-Dependent Classical Dendritic Cells Are Essential for Intestinal T Cell Homeostasis. Immunity 44, 860–874 (2016).

53. Alili, R. et al. Exploring Semi-Quantitative Metagenomic Studies Using Oxford Nanopore Sequencing: A Computational and Experimental Protocol. Genes 12, 1496 (2021).

