## Supplemental figures for "Dietary fibers benefits on glucose homeostasis require type 2 conventional dendritic cells in mice fed a high-fat diet"

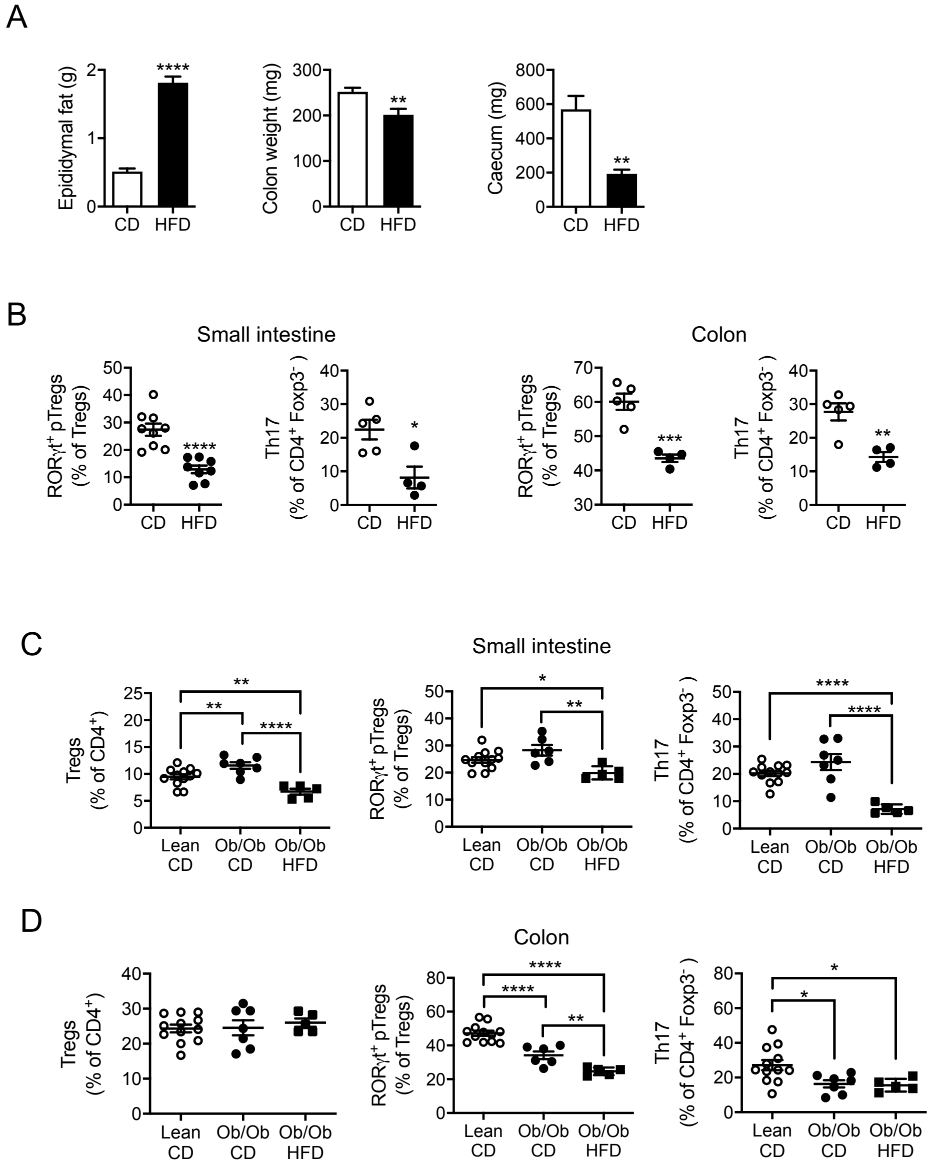


**Figure S1. HFD feeding decreases RORγt⁺ pTreg and Th17 cells in the intestine.**
**(A)** Epidydimal fat mass, colon weight and caecum weight in wild-type mice fed a chow diet (CD) or a high-fat diet (HFD) for 14 weeks. (**B**) Flow cytometry analysis of RORγt⁺ pTregs and Th17 cells in the small intestine and colon of wild-type mice fed a chow diet (CD) or a high-fat diet (HFD) for 14 weeks. (**C**) Flow cytometry analysis of total Tregs, RORγt⁺ pTregs and Th17 cells in the small intestine of control mice (lean) fed a chow diet (CD) and obese *Ob/Ob* mice fed a chow diet (CD) or a high-fat diet (HFD) for 4 weeks. (**D**) Flow cytometry analysis of total Tregs, RORγt⁺ pTregs and Th17 cells in the colon of control mice (lean) fed a chow diet (CD) and obese *Ob/Ob* mice fed a chow diet (CD) or a high-fat diet (HFD) for 4 weeks.

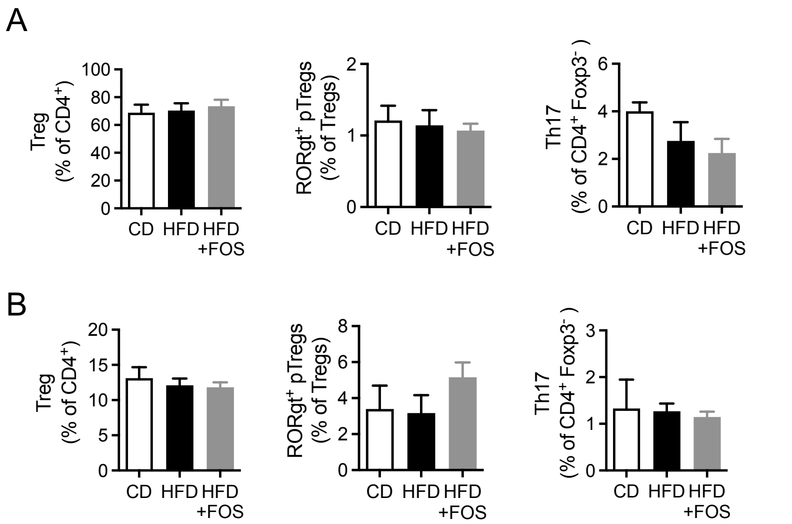


**Figure S2. Dietary fibers do not impact on liver and adipose tissue RORγt+ pTregs and Th17 cells.**

**(A)** Flow cytometry analysis of total Tregs, RORγt+ pTregs and Th17 cells in the epididymal adipose tissue of wild-type mice fed a chow diet (CD), a high-fat diet (HFD) and a high-fat diet supplemented in FOS (HFD+FOS) for 4 weeks. **(B)** Flow cytometry analysis of total Tregs, RORγt+ pTregs and Th17 cells in the liver of wild-type mice fed a chow diet (CD), a high-fat diet (HFD) or a high-fat diet supplemented in FOS (HFD+FOS) for 4 weeks.

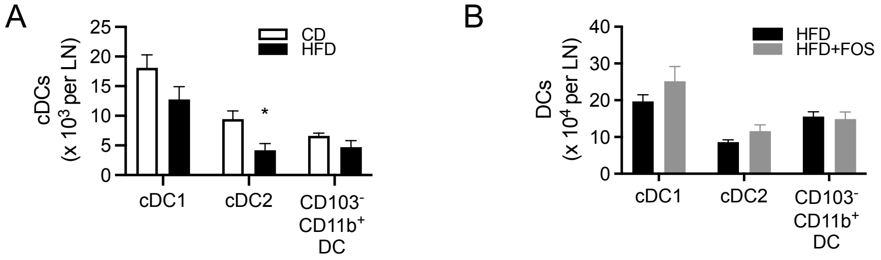


**Figure S3. Impact of the HFD and FOS supplementation on conventional dendritic cell subsets counts.**
**(A)** Conventional dendritic cell (cDC) counts in the mesLNs of wild-type mice fed a chow diet (CD) or a high-fat diet (HFD) for 4 weeks. **(B)** Conventional dendritic cell (cDC) counts in the mesLNs of wild-type mice fed a high-fat diet (HFD) or a high-fat diet supplemented in FOS (HFD+FOS) for 4 weeks.

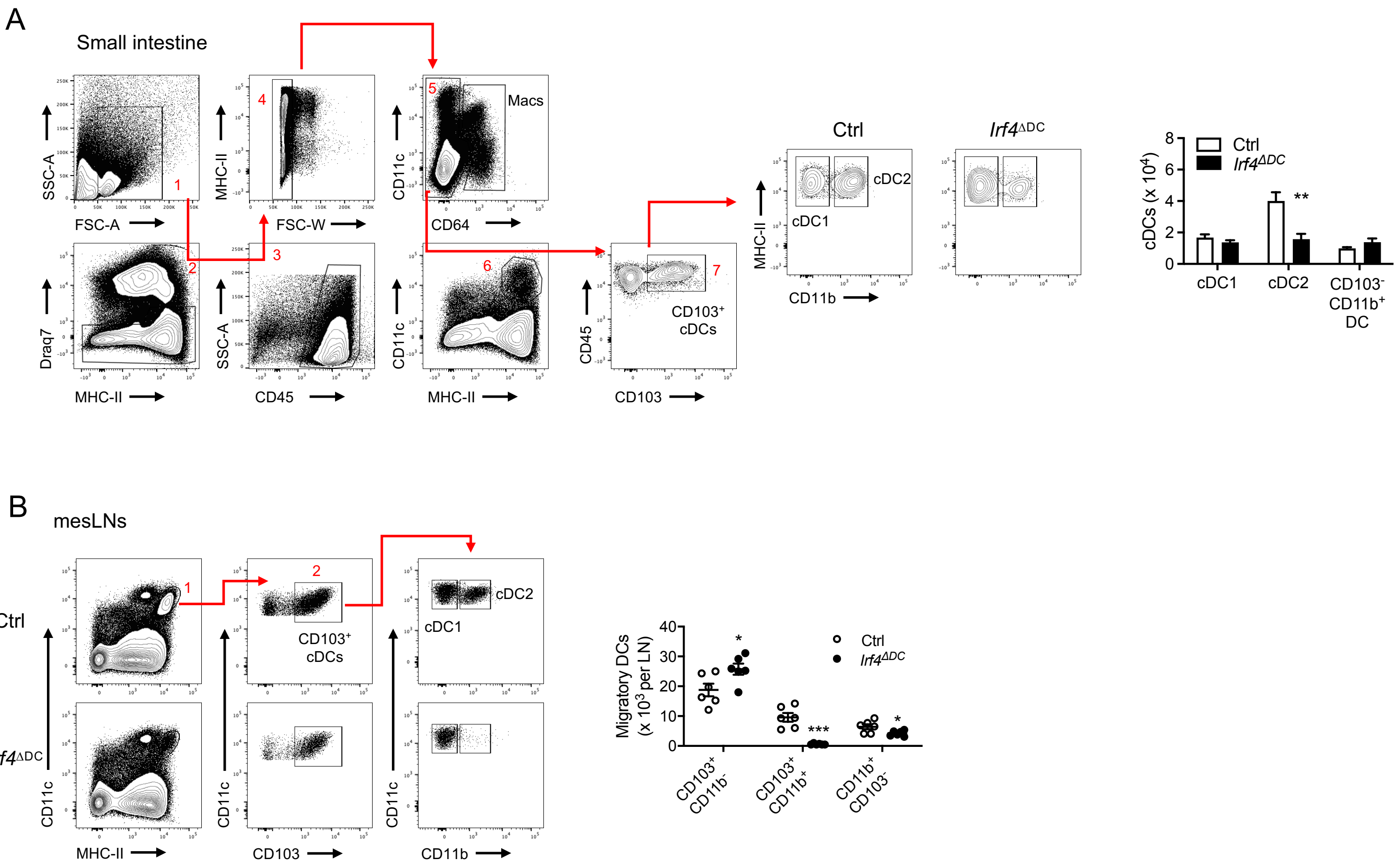


**Figure S4. Dendritic cell subsets in mice invalidated for *Irf4* in CD11c-expressing cells.**
(**A**) Flow cytometry plots and analysis of dendritic cells subsets ($CD103^+$ $CD11b^-$ cDC1 and $CD103^+$ $CD11b^+$ cDC2) in the small intestine of mice lacking *Irf4* in dendritic cells (*Irf4*$^{\Delta DC}$) and *Irf4*$^{flox/flox}$ controls (Ctrl). (**B**) Flow cytometry plots and analysis of dendritic cells subsets ($CD103^+$ $CD11b^-$ cDC1 and $CD103^+$ $CD11b^+$ cDC2) in the mesenteric lymph nodes (mesLNs) of mice lacking *Irf4* in dendritic cells (*Irf4*$^{\Delta DC}$) and *Irf4*$^{flox/flox}$ controls (Ctrl).

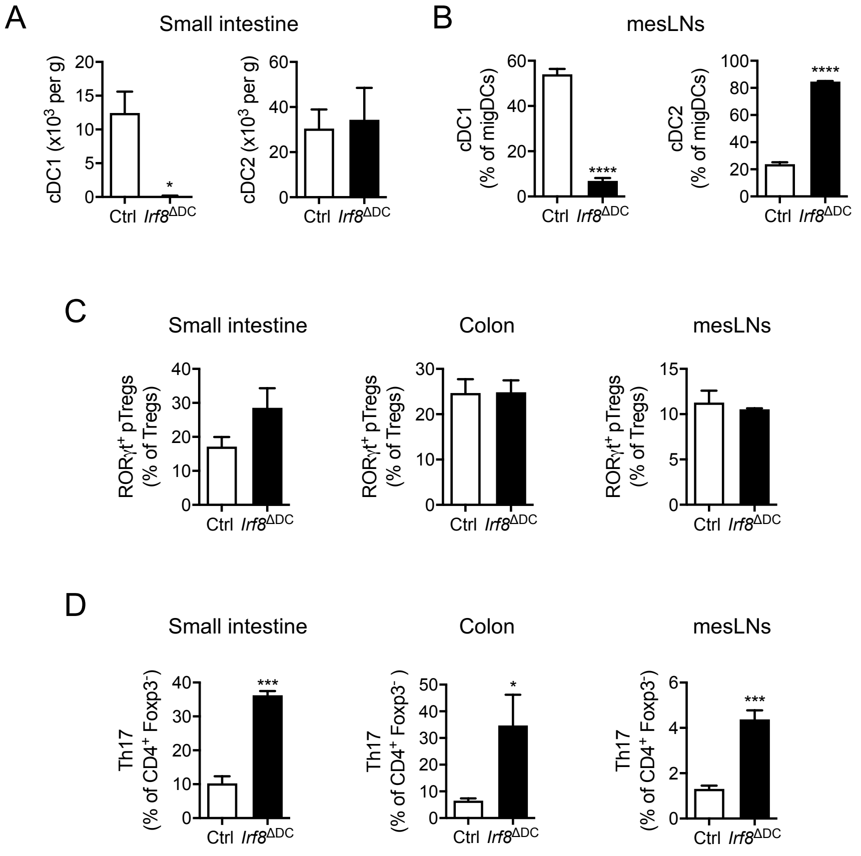


**Figure S5. Analysis of RORγt⁺ pTregs and Th17 cells in mice devoid of cDC1.**
(**A** and **B**) Dendritic cells subsets (CD103$^+$ CD11b$^-$ cDC1 and CD103$^+$ CD11b$^+$ cDC2) in the small intestine (A) and mesenteric lymph nodes (mesLNs) of mice lacking *Irf8* in dendritic cells (*Irf8*$^{\Delta DC}$) and *Irf8*$^{flox/flox}$ controls (Ctrl). (**C** and **D**) Flow cytometry analysis of RORγt$^+$ pTreg (C) and Th17 cells (D) in the small intestine, colon and mesenteric lymph nodes (mesLNs) of mice lacking *Irf8* in dendritic cells (*Irf8*$^{\Delta DC}$) and *Irf8*$^{flox/flox}$ controls (Ctrl).
